## Supplemental File for "Heterogeneous interactions and polymer entropy decide organization and dynamics of chromatin domains"



PACS numbers:


The main paper presents findings of studies on statics and dynamics of a chromatin domain. Through chromatin *in vivo* shows dynamicity and cell-to-cell variability; in literature, the nature of chromatin is mostly quantified through average static properties. In this work, we go beyond the average and quantify the whole phase space by investigating probability distributions and several temporal quantities, and we estimate the timescales of loop formation and stable loop maintenance. The necessary supplementary information for results presented in the main paper is provided here.

#### S1. SIMULATION DETAILS

We consider chromatin as a bead spring chain having optimal intra-chromatin interactions derived from 5C and Hi-C data using an inverse method described below. Taking experimental contact probability data as input, we have computed the optimal interaction strength between different segments of the chromatin using an Inverse Brownian Dynamics (IBD) method that we have developed [1]. We perform Brownian dynamics simulations and compute various static and dynamic quantities as described below. It is essential to include hydrodynamic interactions (HI) while computing dynamic quantities in order to obtain even qualitatively accurate predictions [2–11]. HI accounts for the propagation of velocity perturbations due to the motion of one segment of a polymer chain to other segments through the medium. Several studies that attempt to probe chromatin dynamics have not considered HI. In the present work, we simulate the dynamic nature of chromatin accounting for HI.

##### Governing equations for the bead-spring chain model

The total energy of the chromatin bead spring chain, made up of $N$ beads, is $U = U^{\mathrm{S}} + U^{\mathrm{SDK}}$ where $U^{\mathrm{S}}$ is the spring potential between the adjacent beads $i$ and $(i+1)$, given by

$$U^{\mathrm{S}} = \sum_i \frac{H}{2}(|\boldsymbol{r}_i - \boldsymbol{r}_{i+1}| - r_0)^2 \tag{1}$$

where $\boldsymbol{r}_i$ is the position vector of bead $i$, $r_0$ is the natural length and $H$ is the stiffness of the spring. The Soddemann-Duenweg-Kremer potential ($U^{\mathrm{SDK}}$) is a Lennard-Jones-like potential used to mimic protein-mediated interactions [12, 13] which is given by:

$$U^{\mathrm{SDK}} = \begin{cases} 4\left[\left(\dfrac{\sigma}{r_{ij}}\right)^{12} - \left(\dfrac{\sigma}{r_{ij}}\right)^{6} + \frac{1}{4}\right] - \epsilon_{ij} & r_{ij} \le 2^{\frac{1}{6}}\sigma \\ \frac{1}{2}\epsilon_{ij}\left[\cos\left(\alpha r_{ij}^2 + \beta\right) - 1\right] & 2^{\frac{1}{6}}\sigma \le r_{ij} \le r_c \\ 0 & r_{ij} \ge r_c. \end{cases} \tag{2}$$

Here $r_{ij} = |\boldsymbol{r}_i - \boldsymbol{r}_j|$ is the distance between beads $i$ and $j$ and $\epsilon_{ij}$ is an independent parameter representing the attractive interaction strength between them. $2^{1/6}\sigma$ is the distance at which $U^{\mathrm{SDK}}$ is zero. The cut-off radius of the SDK potential is $r_c = 1.82\sigma$. The SDK potential has the following advantages compared to the Lennard-Jones (LJ) potential:

1. The repulsive part of the SDK potential ($r_{ij} \le 2^{1/6}\sigma$) is constant and remains unaffected by the choice of the parameter $\epsilon_{ij}$.
2. The SDK potential reaches zero at the cut off radius $r_c = 1.82\sigma$, unlike the LJ potential where the energy goes to zero only at infinite distance.
3. As the protein-mediated crosslinking interactions in chromatin are short ranged, the SDK potential models the chromatin crosslinks more appropriately [12, 14].

For the simulation, all the length and time scales are non-dimensionalised with $l_H = \sqrt{k_B T/H}$ and $\lambda_H = \zeta/4H$, respectively where $T$ is the absolute temperature, $k_B$ is the Boltzmann constant, and $\zeta = 6\pi\eta_s a$ is the Stokes friction coefficient of a spherical bead of radius $a$, with $\eta_s$ being the solvent viscosity. The evolution of bead positions in BD simulations is governed by the following Ito stochastic differential equation,

$$\begin{aligned} \boldsymbol{r}_i^*(t^* + \Delta t^*) = \boldsymbol{r}_i^*(t^*) &+ \frac{\Delta t^*}{4}\boldsymbol{D}_{ij} \cdot (\boldsymbol{F}_j^{S*} + \boldsymbol{F}_j^{\mathrm{SDK}*}) \\ &+ \frac{1}{\sqrt{2}}\boldsymbol{B}_{ij} \cdot \Delta\boldsymbol{W}_j \end{aligned} \tag{3}$$

Here $t^* = t/\lambda_H$ is the dimensionless time and $r_i^* = r_i/l_H$ is the dimensionless length. $\Delta \boldsymbol{W}_j$ is a non-dimensional Wiener process with mean zero and variance $\Delta t^*$. The bonded interactions between the beads are represented by a non-dimensional spring force, $\mathbf{F}_j^{S*}$, and the non-dimensional SDK force is $\mathbf{F}_j^{SDK*}$. $\boldsymbol{D}_{ij}$ is the diffusion tensor, defined as $\boldsymbol{D}_{ij} = \delta_{ij}\boldsymbol{\delta} + \boldsymbol{\Omega}_{ij}$, where $\delta_{ij}$ is the Kronecker delta, $\boldsymbol{\delta}$ is the unit tensor, and $\boldsymbol{\Omega}_{ij}$ is the hydrodynamic interaction tensor. We use the regularized Rotne-Prager-Yamakawa (RPY) tensor to compute hydrodynamic interactions (HI) [5],

$$\boldsymbol{\Omega}(\mathbf{r}^*) = \Omega_1 \boldsymbol{\delta} + \Omega_2 \frac{\mathbf{r}^*\mathbf{r}^*}{r^{*2}} \tag{4}$$

for $r^* \geq 2\sqrt{\pi}h^*$

$$\begin{aligned} \Omega_1 &= \frac{3\sqrt{\pi}}{4}\frac{h^*}{r^*}\left(1 + \frac{2\pi}{3}\frac{h^{*2}}{r^{*2}}\right); \\ \Omega_2 &= \frac{3\sqrt{\pi}}{4}\frac{h^*}{r^*}\left(1 - \frac{2\pi}{3}\frac{h^{*2}}{r^{*2}}\right) \end{aligned} \tag{5}$$

and for $r^* \leq 2\sqrt{\pi}h^*$

$$\begin{aligned} \Omega_1 &= 1 - \frac{9}{32}\frac{r^*}{h^*\sqrt{\pi}}; \\ \Omega_2 &= \frac{3}{32}\frac{r^*}{h^*\sqrt{\pi}} \end{aligned} \tag{6}$$

Here $h^*$ is the dimensionless hydrodynamic parameter, defined as $h^* = a/(l_H\sqrt{\pi})$. In the present work, we set $h^* = 0.25$ [10]. All the simulation parameters with their corresponding values are indicated in Table S1.

*Inverse Brownian Dynamics (IBD)*

To simulate chromatin dynamics accurately, one requires the potential energy parameters that specify the interaction strengths ($\epsilon_{ij}$) between different regions of chromatin. We have developed an IBD algorithm that extracts the optimal interaction strengths between bead pairs from an experimental contact probability map [1]. The details of the algorithm are given below:

The essence of the algorithm is to find the average contact probability and iteratively compare it to the reference contact probability obtained from experiments. The average contact probability $p_m$ of the bead-pair $m$ is given by

$$p_m = \langle \hat{p}_m \rangle = \frac{1}{Z}\int d\Gamma\, \hat{p}_m \exp(-\beta\mathcal{H}) \tag{7}$$

Here $Z = \int d\Gamma \exp(-\beta\mathcal{H})$ is the partition function and $\hat{p}_m$ is an indicator function which indicates when contact occurs between the bead pair represented by index $m$. $\hat{p}_m$ is 1 if the distance between the beads is less than the cut-off distance of the indicator function, $r_p^*$, and

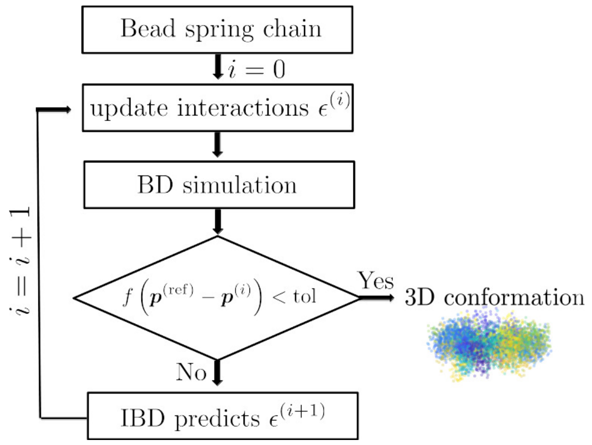


FIG. S1: A flowchart representation of the inverse Brownian dynamics algorithm. Each iteration consists of BD simulation, calculation of contact probability and revision of interaction strength. Here, $p^{(\mathrm{ref})}$ represents the reference contact probability matrix, and $p^{(\mathrm{i})}$ represents the contact probability matrix from simulations at iteration i. This figure is reproduced from Kumari et al. [1].

0 otherwise. For this work $r_p^* = r_c^* = 1.82\sigma$ is taken as the cut-off distance/capture radius of the SDK potential. For our studies $\sigma$ is of the order of 36nm($1.82\sigma \approx 65$nm). We do not know the value of the capture radius ($r_p^*$) precisely. However, Giorgetti et al. [15], have previously systematically varied a similar capture radius parameter and have reported a value of roughly the same order as that used here (between 50nm and 100nm).

In a separate study, we have used the SDK potential to systematically study the behaviour of polymers in a poor solvent, with the capture radius as a parameter, and have reproduced the standard scaling in polymer physics accurately. It was found that $r_c = 1.82\sigma$ was the optimum value for reproducing homogenous polymer behaviour in poor solvents [13].

As the interaction strengths are defined between a pair of beads, we use a single index ($m$) to define a specific bead pair ($i$ and $j$) as $m = \frac{1}{2}[i(i-1)] + [j-(i-1)]$. Here $i$ varies from 2 to $N$, and $j$ varies from 1 to $(i-1)$ for a matrix of size $N$. $\hat{p}_m$ is 1 if the distance between the beads is less than the cut-off distance of the indicator function, $r_c^*$, and 0 otherwise. We intend to target the experimentally obtained contact probability $p_m^{\mathrm{ref}}$ by adjusting the well-depth of SDK attractive interactions $\epsilon_m$. The Taylor series expansion of $\langle \hat{p}_m \rangle$ about the interaction strength $\epsilon_m$ after neglecting higher order terms is

$$\langle \hat{p}_m \rangle(\epsilon_m + \Delta\epsilon_m) = \langle \hat{p}_m \rangle(\epsilon_m) + \sum_n \chi_{mn}\,\Delta\epsilon_n \tag{8}$$

where $\Delta\epsilon_m$ is the change in the interaction strength, and

TABLE S1: Numerical values of all the parameters used in the simulation

| Simulation parameters | symbols | values |
|---|---|---|
| no. of beads | $N$ | 50 |
| cut-off radius | $r_c^*$ | 1.82 $\sigma^*$ |
| natural length of the spring | $r_0^*$ | 1 $\sigma^*$ |
| $U^{\mathrm{SDK}}$ length scale | $\sigma^*$ | 1 |
| hydrodynamic parameter | $h^*$ | 0.25 |
| SDK parameter | $\alpha$ | 1.53063 |
| SDK parameter | $\beta$ | 1.21311 |
| length scale | $l_H$ | 36 nm |
| time scale | $\lambda_H$ | 0.1 s |
| interaction strength | $\epsilon_{ij}$ | from IBD |

the susceptibility matrix

$$\chi_{mn} = \frac{\partial \langle \hat{p}_m \rangle}{\partial \epsilon_n} = \frac{\partial}{\partial \epsilon_n} \left[ \frac{1}{Z} \int d\Gamma \hat{p}_m \exp(-\beta \mathcal{H}) \right] \tag{9}$$

$$\begin{aligned} \chi_{mn} =& \beta \left[ \frac{1}{Z} \int \hat{p}_m b_n \exp(-\beta \mathcal{H}) d\Gamma \right. \\ & \left. - \langle \hat{p}_m \rangle \frac{1}{Z} \int b_n \exp(-\beta \mathcal{H}) d\Gamma \right] \end{aligned} \tag{10}$$

$$\chi_{mn} = \beta \left[ \langle \hat{p}_m b_n \rangle - \langle \hat{p}_m \rangle \langle b_n \rangle \right]$$

where

$$b_n = -\frac{\partial \mathcal{H}}{\partial \epsilon_n} \tag{11}$$

Replacing the left hand side of Eq. 8 with the target contact probability $p_m^{\mathrm{ref}}$ obtained from experiment, we get

$$p_m^{\mathrm{ref}} - \langle \hat{p}_m \rangle = \sum_n \chi_{mn} \Delta \epsilon_n \tag{12}$$

Equation 12 can be solved for any particular iteration step as

$$\epsilon_n^{(i+1)} = \epsilon_n^{(i)} + \lambda \sum_m C_{nm}^{(i)} \left( p^{\mathrm{ref}} - \langle \hat{p}_m \rangle^{(i)} \right) \tag{13}$$

where the matrix $\mathsf{C}$ is the *pseudo-inverse* of the matrix $\chi$ [1], superscript $i$ represents the iteration number, $\lambda$ denotes the damping factor with $0 < \lambda < 1$, and $\epsilon_n^{(i+1)}$ is the well-depth of the SDK attractive interaction for the next iteration step.

The flowchart of the IBD methodology is given schematically in Fig. S1. We start with the initial guess values of interaction strengths, simulate the polymer following the conventional forward Brownian dynamics method and obtain the simulated contact probabilities in the steady state. The interaction strengths are revised for the next iteration, depending upon the difference between the simulated and the known experimental contact probabilities. We perform several iterations of the loop (i.e., BD simulation, calculation of contact probability, revision of interaction strength) until the error between the simulated and experimental contact probabilities is less than a predetermined tolerance value. Using these optimal interaction strengths, we study the dynamics of the chromatin domain. In the present work, we have applied the IBD technique to the two chromatin domains ($\alpha$-globin gene locus and a domain in Chr7) and investigate the static and dynamic properties of the chromatin domain. The error between the reference contact map and the contact map computed from simulations decreases and converges to a small value, smaller than a desired tolerance value. The optimised interaction strength ($\epsilon_{ij}$) values for $\alpha$-globin are shown in the Fig. S2.

### S2. EQUATION PREDICTING INTERACTION STRENGTH ($\epsilon_{ij}$) FROM CONTACT PROBABILITY ($p_{ij}$)

To understand how $\epsilon_{ij}$ depends on $p_{ij}$ and the genomic separation ($|i-j|$), we plotted the data from IBD as shown in Fig. S3(a). This gave us a hint that the $\epsilon_{ij}$ increases by $p_{ij}$ logarithmically ($\epsilon_{ij}$ increases with a factor of 1 while $p_{ij}$ increases by a factor of 10). We also observed that for every $|i-j|$, there exist a minimum value of $p_{ij}$ which we define as $p_{\min}(i-j)$. The $p_{\min}(i-j)$ for every $|i-j|$ can be easily obtained from the contact probability matrix. This gave us the formula described in the Eq. 1 of the main text. Here, we discuss how well equation. 1 in the main text predicts $\epsilon_{ij}$ values. Fig. S3(b) shows the relation between interaction strength and contact probability from the IBD simulation and Eq.1 of the main manuscript for the Chr.7 region from the IMR90 cell line. A simple way to test the equation is to have a scatter plot $\epsilon_{ij}$ from IBD in the X-axis and $\epsilon_{ij}$ from the equation (Eq. 1 in the main text) in the Y-axis. as shown in Fig. S3(c). They fall near the line $y = x$, implying a good correlation between the two. This formula would

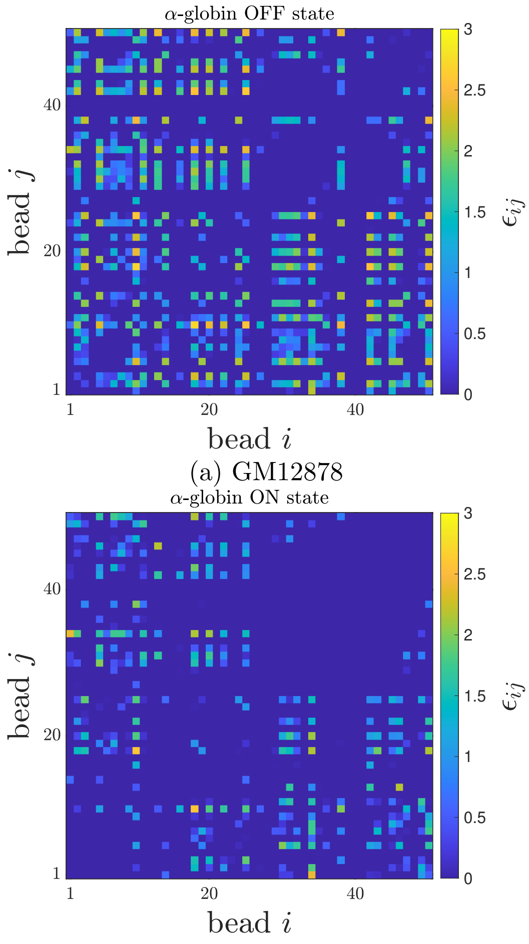


(a) GM12878

(b)K562

FIG. S2: (a) and (b) show the optimised interaction strength for OFF (GM12878) and ON (K562) state of $\alpha$-globin, respectively.

immensely help in generating excellent initial guess values of $\epsilon_{ij}$ and will lead to faster convergence of the IBD.

A naive guess for such a formula would have been $\epsilon_{ij} \propto -\log(p_{ij})$. In Fig. S3(d), we are testing this formula and found that this simple relation does not hold.

### S3. SCALING OF 3D DISTANCE WITH GENOMIC DISTANCE: COMPARISON WITH EXPERIMENTS

We computed the spatial distance in the repressed state of human $\alpha$-globin gene (OFF state) and found $\langle r^*_{ij}\rangle \sim s^{\nu}_{ij}$ with a scaling exponent $\nu = 0.38$ (Fig. S4). This suggests near close packing within the chromatin domain (red symbols). As a "control", we also simulated a self-avoiding walk (SAW) polymer with no attractive interaction ($\epsilon_{ij} = 0$) which results in $\nu = 0.6$ as expected (black symbols) [17]. Our chromatin model predicts a spread (variability) in the 3D distance (red symbols) which is absent in the control revealing the implications of heterogeneous intra-chromatin interactions. A recent microscopy study [16] on a mouse ESC chromatin domain also showed a similar behaviour – both the scaling (slope) and the variability in the experimental and simulation data are comparable without any fitting parameter. This is a validation that macroscopic polymer properties of the chromatin domain in our simulation accurately represent what is observed in realistic systems. The $y$-intercept of the experimental data gives us the size ($\sigma = l_H$) of the 10kb chromatin (a single bead in our simulation). For this experimental system (mouse chromosome 6, 1.2MB in Szabo et al. [16]) we get $l_H = 22$nm. Even though we do not have such extensive spatial distance data (between all pairs) for human $\alpha$ globin, we used the available FISH data for the distance between a single pair of $\alpha$ globin segments and deduced the $l_H = 36$nm. Throughout this paper, $l_H = 36$nm and $\lambda_H = 0.1$s are used to convert all non-dimensional lengths and times into standard units, and we will present quantities in both units. The reasons for the choice of both these specific values are discussed in greater detail in the next section.

### S4. CONVERSION OF NON-DIMENSIONAL LENGTH AND TIME TO STANDARD UNITS

In our simulations, all quantities are computed in dimensionless units as described earlier. To convert these dimensionless numbers to standard units having appropriate dimensions, we need to determine a length scale and a timescale. By comparing our simulations with appropriate experimental observations, we deduce values of characteristic length and time scales that can be used for the unit conversion as follows:

**Length scale:** Even though the 3D distances between the genomic segments of $\alpha-$globin are not available, the 2D distance between the two probes located at $34,512$ - $77,058$ bp and $386,139$ - $425,502$ bp for GM12878 (OFF state) was found to be $318.8 \pm 17.0$ nm from the 2D FISH [18]. We computed the average 2D distance ($= 8.81$) between the corresponding bead pair (bead 5 and bead 40) from our simulation by averaging it over all the three 2D planes ($xy$, $yz$, $zx$) as depicted in Fig. S5. By comparing 2D distance values obtained from simulation and experiment, we estimate the characteristic lengthscale in our simulation as $l_H = 318.8/8.81 \approx 36$nm. We use this value of $l_H$ to convert all non-dimensional lengths to standard units.

**Time scale:** The timescale in our simulation is given by:

$$\lambda_H = \frac{\zeta}{4H} = \frac{6\pi\eta_s a^3}{4k_\mathrm{B}T} \tag{14}$$

where $H$ is the spring constant, $T$ is the absolute temperature, $k_B$ is the Boltzmann constant, and $\zeta = 6\pi\eta_s a$ is the Stokes friction coefficient of a spherical bead of

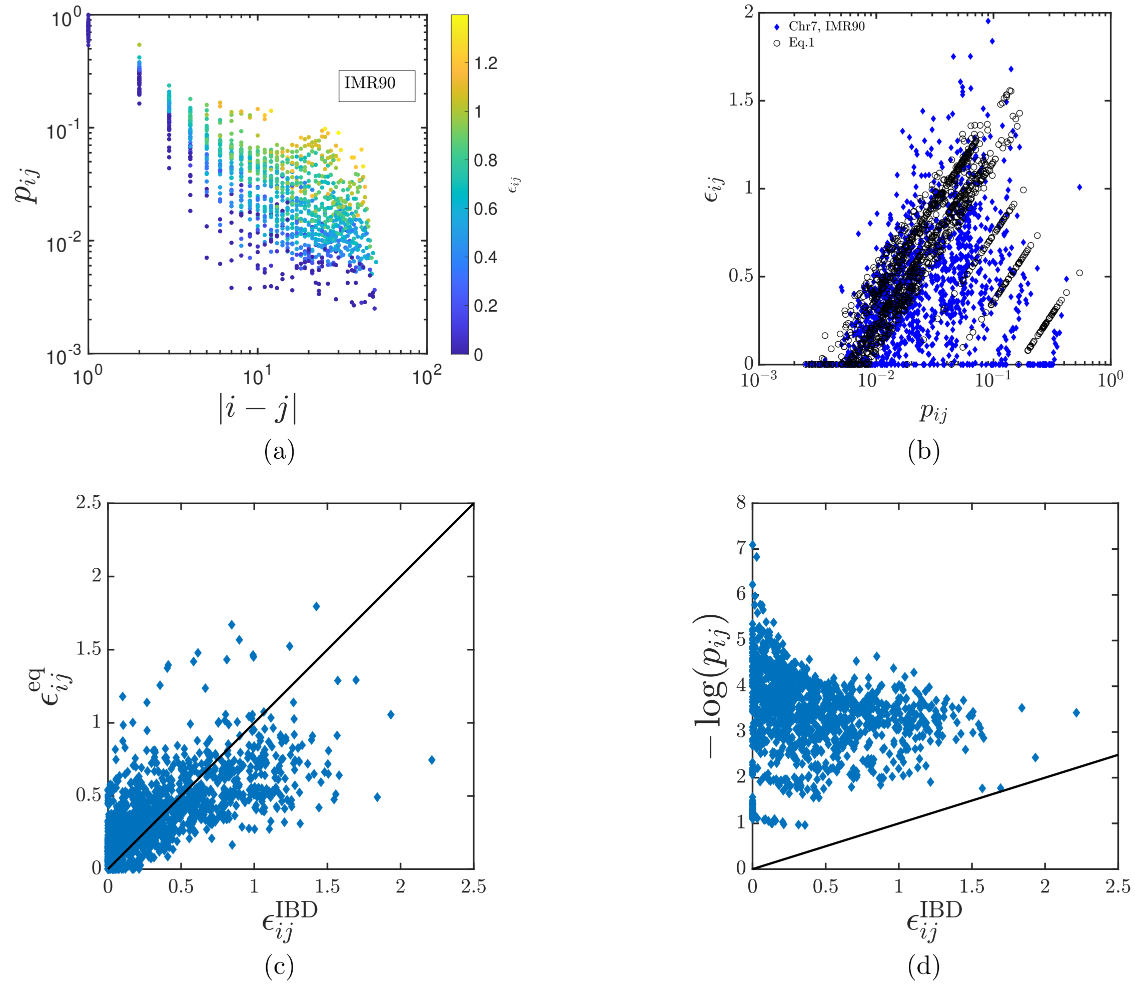


FIG. S3: (a) Our prediction of the relation between interaction strengths ($\epsilon_{ij}$), contact probability and segment length ($|i-j|$) from IBD simulations. See colorbar for $\epsilon_{ij}$. (b) Prediction of the relation between interaction strength and contact probability from the IBD simulation (filled diamonds) and Eq.1(black open circles) of the main manuscript for the Chr.7 region from IMR90 cell line. (c) Scatter plot of $\epsilon_{ij}$ from IBD in the $x$-axis and $\epsilon_{ij}$ from the equation (Eq. 1 in the main text) in the $y$-axis. There is a good positive correlation between the two, and the points fall near the line $y = x$ suggesting that the equation gives a reasonable estimate of the $\epsilon_{ij}$. (d) Scatter plot of $\epsilon_{ij}$ from IBD in the $x$-axis and the negative of $\log p_{ij}$ in the $y$-axis. $\epsilon_{ij} = -\log p_{ij}$ is a naive guess for the interaction strengths. But the points highly deviating from the $y = x$ line suggest that $-\log p_{ij}$ is not a good estimate of $\epsilon_{ij}$. Both the plots here are for K562 cell type.

radius $a$ where $\eta_s$ is the solvent viscosity. For our problem $a = h^* l_H \sqrt{\pi} \approx$ 16nm. However, we do not know the precise viscosity of the solvent in the nucleus. There are many estimates ranging over several orders of magnitude from $10^{-3}$ Pa.s to $10^3$ Pa.s. [19, 20]. Given this degree of variability, we decided to use a simple method to estimate time, based on recent experimental reports of chromatin dynamics. Chromatin segments under microscope seems to "diffuse" around in a region having the size of the order of $\approx 0.1(\mu m)^2$ within a timescale of $\approx$ 50 seconds [21]. This leads to a diffusion coefficient ($D$) of the order of 500 nm$^2$/s, and a timescale

$$\lambda_H = \frac{a^2}{4D} = \frac{(16\ \text{nm})^2}{4 \times 500\ \text{nm}^2/\text{s}} = 0.12\ \text{s} \qquad (15)$$

Since the calculation is to estimate the order of magnitude number, throughout this work, we use $\lambda_H = 0.1$ s. Interestingly, this also corresponds to an effective viscosity roughly in the middle of the wide range estimated previously. These values of $l_H$ and $\lambda_H$ are considered through out the manuscript unless stated otherwise.

### S5. DISTANCE PROBABILITY DISTRIBUTIONS

The analytical expression given by des Cloizeaux [22] for the distance probability distribution for a self-

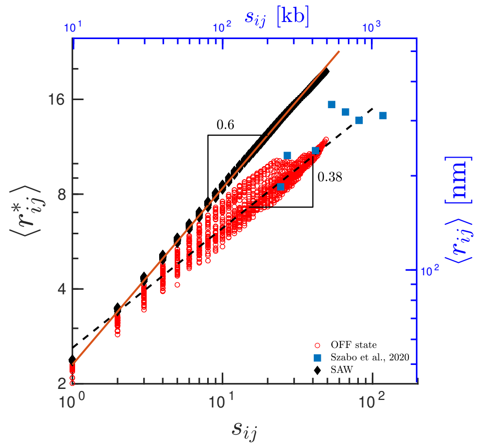


FIG. S4: Average 3D distances between all bead-pairs as a function of corresponding genomic distances for the control simulations (SAW, black symbols), chromatin domain that we simulated (red symbols) and comparison with experimental data from [16] (blue symbols). The major axes (lower $x$ and left $y$) represent quantities in dimensionless units (see methods) while the other axes (upper $x$ and right $y$) represent the same in standard units.

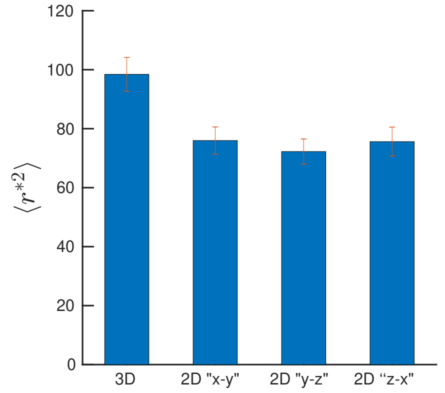


FIG. S5: Depiction of the calculation of 2D distance between bead 5 and bead 40 from simulation.

avoiding walk (SAW) polymer is

$$p(r^*) = C[r^*]^{\theta+2} e^{-[Kr^*]^{\frac{1}{1-\nu}}} \tag{16}$$

Here $\nu$ is the Flory exponent, $\theta$ is a geometrical exponent and the coefficients $C$ and $K$ are given by

$$K^2 = \frac{\Gamma([\theta+d+2][1-\nu])}{\Gamma([\theta+d][1-\nu])}$$

$$C = 4\pi \frac{[\Gamma([\theta+d+2][1-\nu])]^{\frac{\theta+d}{2}}}{[\Gamma([\theta+d][1-\nu])]^{\frac{\theta+d+2}{2}}}$$

where $d$ is the dimension. Since our simulations are in 3D, $d = 3$. The geometrical exponent $\theta$ takes different values in the following three cases

1. Case 1: When both beads are the end beads of the polymer ($\theta = \theta_0$),

2. Case 2: When one of the beads is at the end and the other bead is an intermediate bead within the chain ($\theta = \theta_1$),

3. Case 3: When both the beads are intermediate beads ($\theta = \theta_2$)

As the coefficients $C$ and $K$ depend on $\theta$, they take different values in each of the above cases. Following the findings of des Cloizeaux [22], Witten and Prentis [23], Duplantier [24] and Hsu et al. [25], one can determine that $\theta_0 = 0.267$, $\theta_1 = 0.461$ and $\theta_2 = 0.814$ [26]. Simulating a SAW polymer, we compared the probability distribution for all the three cases with the corresponding analytical expressions using the appropriate values of $\theta$. Fig. S6 show the validation for case 1 and 2, while the validation for case 3 has been shown in the main text. As can be seen, the simulations are in excellent agreement with the analytical expression. To the best of our knowledge, this is the first comparison of exact numerical results with the analytical expression proposed by des Cloizeaux.

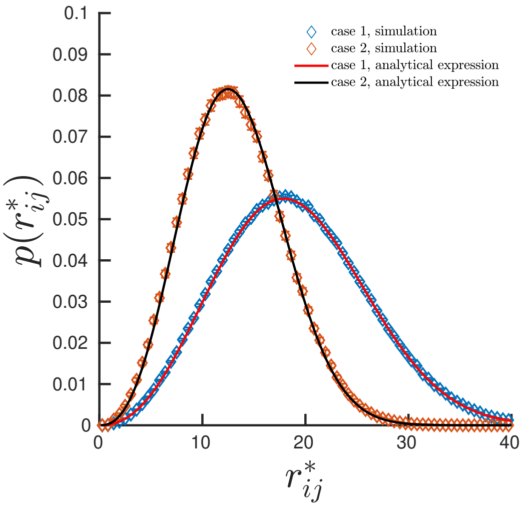


FIG. S6: Comparison of distance probability distributions obtained from the simulation of a SAW chain with the analytical expression for case 1 - where both beads are end beads, and case 2 - where one bead is at the chain end and the other bead is an intermediate bead.

We studied the distance probability distributions of various bead pairs, revealing a distribution with two peaks where one of the peaks is dominated by the entropy of the polymer (genomic separation). The other peak emerges with an increase in the interaction strength. Fig. S7(a) depicts the Cumulative distance distribution $C(r^*)$ for various bead-pairs at same genomic separation ($s_{ij} = 25$) experiencing different interaction strengths. Differences in these plots can be easily noticed only at the small $r_{ij}$ while they look similar overall (see inset). The

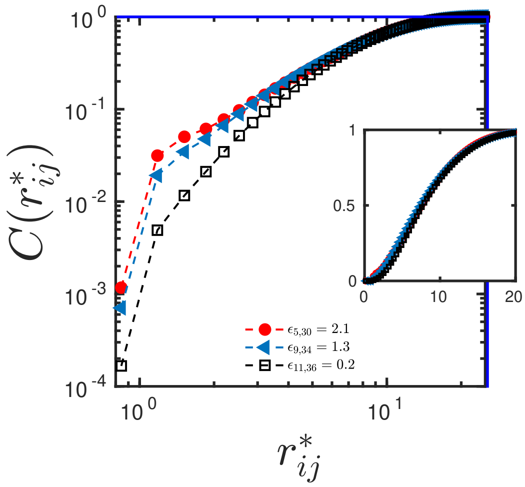


(a)

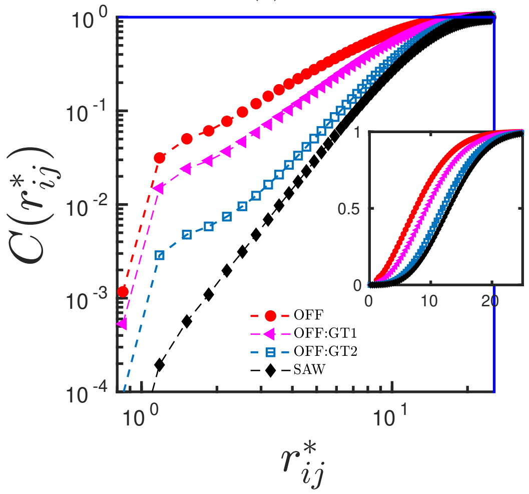


(b)

FIG. S7: (a) Cumulative distance distribution $C(r^*_{ij})$ for various bead-pair with the same genomic separation $s_{ij} = 25$ in the OFF state of $\alpha$-globin gene in log-log scale. The same is indicated in linear scale in the inset. (b) Comparison of $C(r^*_{ij})$ for the chromatin domain under different "epigenetic states" (see text).

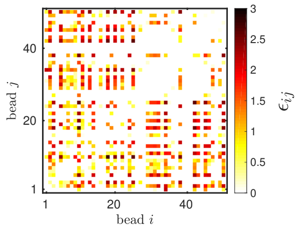


(a) OFF state (GM12878)

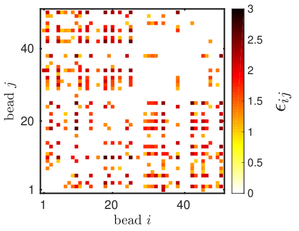


(b) OFF:GT1

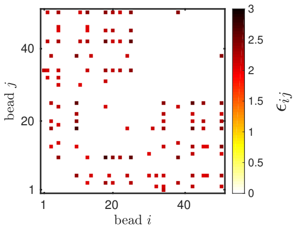


(c) OFF:GT2

FIG. S8: (a) The interaction strength obtained form the IBD for GM12878 $\alpha$-globin gene. (b) A subset of (a) where only the interaction strengths greater than $1k_{\mathrm{B}}T$ are considered. (c) A subset of (a) where the interaction strengths greater than $2k_{\mathrm{B}}T$ are considered.

same has been depicted for a specific bead-pair $(5, 30)$ in different epigenetic states in Fig. S7(b). The difference, in this case, is not only observable for small $r_{ij}$, but for the whole regime of $r_{ij}$ as can be seen in the inset.

### S6. INTERACTION STRENGTH OF PERTURBED STATES

The changes introduced into the pair-wise chromatin interactions in the simulations can be considered as equivalent to epigenetic changes. In this spirit, we have perturbed the interaction strengths systematically to model different epigenetic-like states. The four major states considered in this work are as follows:

1. OFF - all interactions obtained through the IBD for $\alpha$-globin gene locus in GM12878 (off/repressed state) cell line are considered. These values are obtained in our prior work [1] and are displayed in Fig. S8(a) with a heatmap representation.

2. OFF:GT1 - only interactions above $1k_BT$ are considered (see Fig. S8(b)). All other interactions ($< 1k_BT$) are taken as $\epsilon_{ij} = 0$ (i.e., there is only steric hindrance between these bead-pairs). This is a subset of the OFF state mentioned in 1.

3. OFF:GT2 - only strong interactions above $2k_BT$ are considered (see Fig. S8(c)). All interactions below $2k_BT$ are taken as $\epsilon_{ij} = 0$.

4. SAW - a polymer with only steric hindrance, as a control. In other words, all attractive interactions are switched off.

Interestingly, Fig. S8(a) and (b) do not look significantly different to the eye, and even in terms of magnitude, only interactions of the order of thermal fluctuation are switched off, yet as demonstrated in the main text, this causes a qualitative change to predictions.

### S7. SIZE AND SHAPE ANALYSIS

Here we discuss various quantities that are used to characterise a chromatin domain. The radius of gyration of the chain, $R_{\mathrm{g}} \equiv \sqrt{\langle R^2_{\mathrm{g}} \rangle}$, where $\langle R^2_{\mathrm{g}} \rangle$ is defined by

$$\langle R^2_{\mathrm{g}} \rangle = \langle \lambda^2_1 \rangle + \langle \lambda^2_2 \rangle + \langle \lambda^2_3 \rangle \tag{17}$$

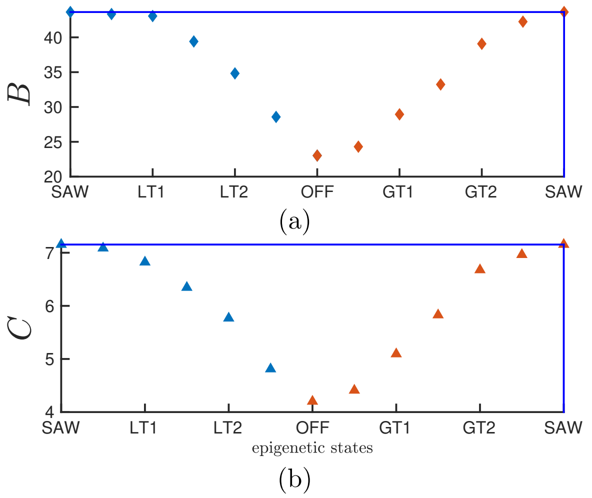


FIG. S9: (a) and (b) represent the asphericity and acylindricity, respectively. $x$-axis represents different interaction states with extreme ends representing the control (SAW) polymer and the OFF state (WT) in the middle. LT1 (LT$x$) indicate that all interactions below $1k_BT$ ($xk_BT$)are present in the polymer. Similarly GT1 (GT$x$) indicate that all interactions above $1k_BT$ ($xk_BT$) are present.

with, $\lambda_1^2$, $\lambda_2^2$, and $\lambda_3^2$ being the eigenvalues of the gyration tensor $\mathbf{G}$ (arranged in ascending order), with

$$\mathbf{G} = \frac{1}{2N^2}\sum_{i=1}^{N}\sum_{j=1}^{N}\mathbf{r}_{ij}\mathbf{r}_{ij} \tag{18}$$

Note that, $\mathbf{G}$, $\lambda_1^2$, $\lambda_2^2$, and $\lambda_3^2$ are calculated for each trajectory in the simulation before the ensemble averages are evaluated [14, 27–32]. We examined a large ensemble of configurations and computed shape properties such as the asphericity ($B$), the acylindricity ($C$), the degree of prolateness ($S$), and the shape anisotropy ($\kappa^2$), as defined below,

$$B = \langle\lambda_3^2\rangle - \frac{1}{2}\left[\langle\lambda_1^2\rangle + \langle\lambda_2^2\rangle\right] \tag{19}$$

$$C = \langle\lambda_2^2\rangle - \langle\lambda_1^2\rangle \tag{20}$$

$$S = \frac{\left\langle(3\lambda_1^2 - I_1)(3\lambda_2^2 - I_1)(3\lambda_3^2 - I_1)\right\rangle}{\left\langle(I_1)^3\right\rangle} \tag{21}$$

$$\kappa^2 = 1 - 3\frac{\langle I_2\rangle}{\langle I_1^2\rangle} \tag{22}$$

Fig. S9(a) and (b) show the asphericity ($B$) and the acylindricity ($C$) for a chromatin domain for various epigenetic states. Both $B$ and $C$ increases monotonically as we go from the OFF state to a SAW.

TABLE S2: The values for the shape quantities and $R_g^2$ for the GT and LT case of $\alpha$-globin gene. This data is plotted in Fig.2 of the main manuscript.

| | LT | | | GT | | |
|---|---|---|---|---|---|---|
| $\epsilon$ | $R_g^2$ | $B/R_g^2$ | $C/R_g^2$ | $R_g^2$ | $B/R_g^2$ | $C/R_g^2$ |
| 0 | 66.3 | 0.658 | 0.107 | 37.6 | 0.612 | 0.183 |
| 0.5 | 66.1 | 0.655 | 0.106 | 39.3 | 0.618 | 0.181 |
| 1 | 66 | 0.652 | 0.103 | 45.8 | 0.631 | 0.173 |
| 1.5 | 60.6 | 0.650 | 0.104 | 51.9 | 0.640 | 0.175 |
| 2 | 54.4 | 0.640 | 0.105 | 59.9 | 0.651 | 0.169 |
| 2.5 | 45.4 | 0.628 | 0.106 | 64.1 | 0.658 | 0.164 |
| 3 | 37.6 | 0.612 | 0.112 | 65.8 | 0.663 | 0.163 |

### S8. STEP-WISE CONVERGENCE OF IBD

To classify the contact probabilities into peaks of various strengths, we adopted the following strategy, which we call "peak detection". Given a segment length, we first calculate the average ($\bar{p}_{|i-j|}$) and standard deviation ($\sigma_{|i-j|}$) of contact probabilities. We then identified the individual contact probability of the bead pair as a prominent peak if the contact probability is greater than the summation of average and standard deviation of contact probability at that segment length ($p_{ij} > \bar{p}_{|i-j|} + \sigma_{|i-j|}$). Similarly, we identified the intermediate peaks if the contact probability is greater than the average contact probability at that segment length ($p_{ij} > \bar{p}_{|i-j|}$). For the stepwise optimization of IBD, we first optimized only for the prominent peaks. Once the prominent peaks are recovered, we optimized for the intermediate peaks. In the end, we optimize for the full contact probability matrix. For instance, Fig. S10 shows the stepwise IBD optimization of the domain in the K562 cell line. Similar to the case for IMR90 discussed in the main text, here, the first row is for the prominent peaks showing the reference contact probability on top, optimized interaction strength at the middle and recovered contact probability at the bottom. Similarly, the second and third column is for the intermediate and full contact probability matrix. Fig. S11(a) and (b) show the optimised interaction strength for the IMR90 and K562 cell line. The step-wise convergence process makes the convergence of the whole domain easier; i.e., it is easier to obtain the correct interaction strength values that would reconstruct the experimentally observed contact maps. Given that the landscape is complex with possible metastable states/barriers, it is not guaranteed that the system will converge to a HiC-like state within a reasonable simulation time. A naive effort to converge accounting for contacts of all bead-pairs often get stuck in states with large error values and fails to converge.

After the convergence of the IBD, we simulated the system with optimal interaction strengths and investigated the relation between the contact probability and the spatial distance between all segment pairs of chromatin. Fig. S12 shows our prediction of the mean 3D distance between every pair of beads ($r_{ij}$) as a function of their corresponding contact probability ($p_{ij}$) for cell line K562. One can observe that there is a broad distribution of 3D distances around the mean value.

### S9. QUANTIFYING THE TEMPORAL NATURE OF CHROMATIN DOMAINS

To study the dynamics of the chromatin domain, we quantified several temporal quantities such as relaxation time, loop formation time and contact time. Temporal variation in properties of chromatin can be found by observing the spatial distance between two segments as a function of time.

As a first step towards quantify dynamics, we extracted the longest relaxation time from the end-to-end vector autocorrelation function $\langle \mathbf{R}^*_{\mathrm{E}}(0) \cdot \mathbf{R}^*_{\mathrm{E}}(t^*) \rangle / \langle \mathbf{R}^{*2}_{\mathrm{E}}(0) \rangle$ where $\mathbf{R}^*_{\mathrm{E}} = |r^*_1 - r^*_{50}|$. This computes the time-dependent correlation of the end-to-end vector. What it essentially measures is how long does it take for the end-to-end beads to fluctuate around its equilibrium value. Fig. S13 displays the autocorrelation function decay with time not accounting for HI. The main paper shows the same with HI and the extracted relaxation time for both (with and without HI). In the OFF state, intra-chromatin interactions are high compared to the SAW; as can be seen from the $p(r^*)$ (Fig.1(c)), in the OFF state, there is a sharp peak in $p(r^*)$ (equivalently there is a steeper minima in free energy, inset of Fig.6(c)) suggesting that the end-to-end beads cannot fluctuate too much. Our analysis of the effective stiffness (Fig.6(c)) further shows that, in the OFF state, the end-to-end points can be imagined as two beads connected by a stiff effective spring (arising from interactions). The spring in the OFF state is stiffer compared to the weaker entropic spring in the SAW. Two points connected by a stiffer spring will have a lower fluctuation time scale (inversely proportional to the effective stiffness). Hence we get a lower timescale in our calculation. Another way of thinking is that the relaxation time will be proportional to $R_g^2$. For the OFF state, $R_g$ is small leading to a smaller relaxation time.

We then looked at the loop formation time ($t^*_{\mathrm{L}}$) defined as the time taken for a pair of beads to meet ($r^*_{ij} < r^*_{\mathrm{C}}$) for the first time, starting from a random equilibrium configuration. Fig. S14 showcases the spatial distance between a specific bead pair (bead 5 and bead 30) with time. It can be easily observed that within a single trajectory, bead-pairs come in contact several times. We also looked at the parameters affecting average loop formation time ($\langle t^*_{\mathrm{L}} \rangle$). Fig. S15(a) and (b) depict $\langle t^*_{\mathrm{L}} \rangle$ with genomic separation ($s$) and interaction strength ($\epsilon$), respectively. All the filled symbols in Fig. S15 represent results from simulation, including HI, while the empty symbols represent no-HI case. The study shows that while the HI is crucial in determining the relaxation time of the whole polymer, it does not affect temporal quantities such as $\langle t^*_{\mathrm{L}} \rangle$.

We also examined the distribution of $t^*_{\mathrm{L}}$ which shows a power law nature with exponential tails (see Fig. S16(a) and (b)). The exponential tail is a typical finite size effect. Since chromatin is of finite size, this is expected.

#### Relation between contact time and interaction energy

Fig.8(b) in the main text shows a relation between contact time and interaction energy as $t_c \propto \exp(\epsilon/2)$. However, one would naively expect that the inverse of the time – the rate of breaking the contact – would be proportional to $e^{-\epsilon}$ from the perspective of a simple Kramers' problem. However, note that this expectation is true only if we assume a Kramers' problem for a single particle in a simple potential well with a barrier of height $\epsilon$. Here, even though the $\epsilon$ is the interaction energy between two beads, there is a whole polymer influencing both binding and dissociation. Even if we assume a simple potential well, the only thermodynamic constraint is that the ratio of the binding and dissociation rates should be $e^{\Delta G}$, where $\Delta G$ has contributions from both energy and entropy. Here $\epsilon$ is just the energy part. The ratio can also have an entropy contribution. Hence, it may deviate from $e^{-\epsilon}$

From another point of view, the true energy landscape need not be a simple one. We do not know the precise (multi-dimensional) free energy landscape; hence, individually, the rates (times) could have a factor of 2 (or any factor for that matter which may not be predictable apriori).

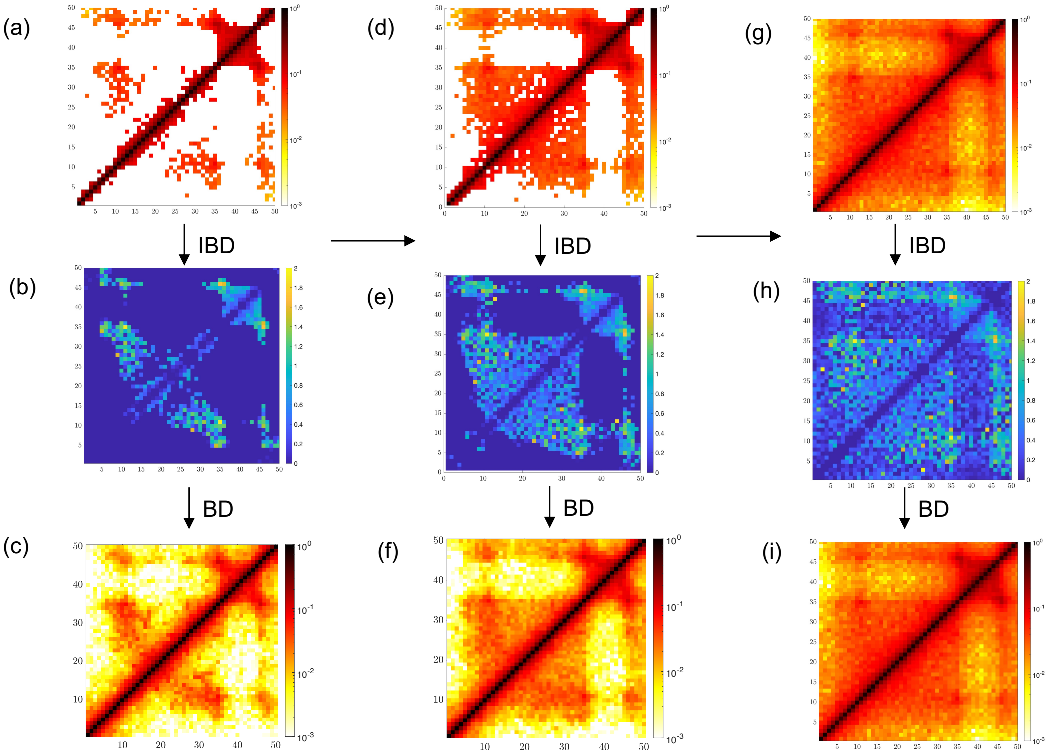


FIG. S10: **Step-wise IBD showing the importance of weak interactions:** Stepwise IBD process for the Chr7 region in the K562 cell line. The IBD optimisation is presented in 3 steps. (I) First column: input is only the prominent probability values (higher than one standard deviation from the mean. i.e., $p_{ij} > \bar{p}_{|i-j|} + \sigma_{|i-j|}$) show in (a) based on the peak detection algorithm. (b) The corresponding optimised $\epsilon_{ij}$ values, and (c) the recovered contact probabilities. (II) Second column: after the first optimisation step, all peaks above the average probability values (shown in (d)) are fed as input (i.e., $p_{ij} > \bar{p}_{|i-j|}$). The corresponding optimised interaction strengths in (e) are improved with some weak interactions appearing in this 2nd step. However note that the recovered contact probability simulated with prominent interactions in (f) is not comparable to the full contact probability in (g). (III) Third column: the complete Hi-C matrix in (g) is fed as input. At the end of this third step the whole contact probability matrix was recovered in (i), and the corresponding optimal interaction strengths are predicted in (h).

[17] R. Bird, C. Curtiss, R. Armstrong, and O. Hassager, Dynamics of polymeric liquids, kinetic theory (volume 2) (1987).

[18] D. Baù, A. Sanyal, B. R. Lajoie, E. Capriotti, M. Byron, J. B. Lawrence, J. Dekker, and M. A. Marti-Renom, Nat. Struct. Mol. Biol. **18**, 107 (2011).

[19] F. Erdel, M. Baum, and K. Rippe, J. Phys.: Condens.

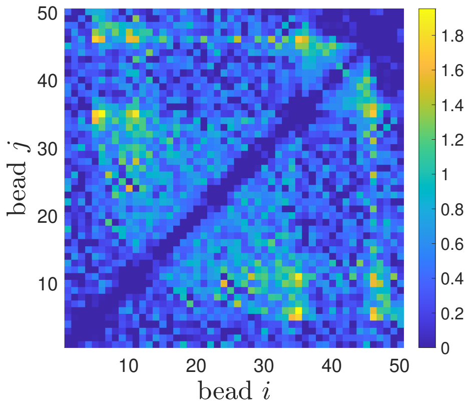


(a) IMR90

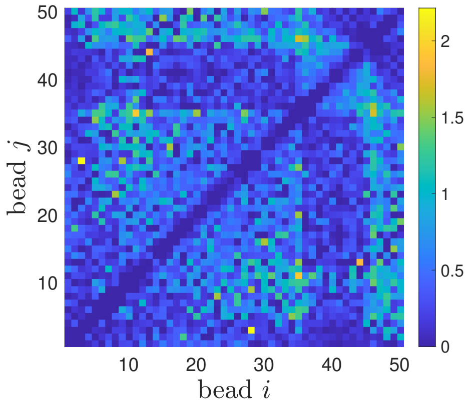


(b)K562

FIG. S11: (a) and (b) show the optimised interaction strength for a domain in IMR90 and K562 cell line, respectively.

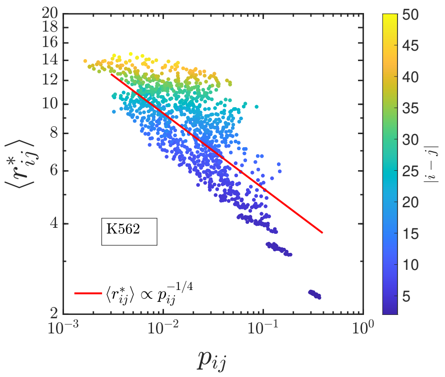


FIG. S12: Our prediction of the mean 3D distance between every pair of beads ($r_{ij}$) as a function of their corresponding contact probability ($p_{ij}$) for cell line K562. The color indicates genomic distance $|i-j|$ between the pair of beads (in units of bead size, see sidebar). Curve with a power-law relation $r_{ij} \propto p_{ij}^{(-1/4)}$ is plotted (red line) as a guide to the eye.

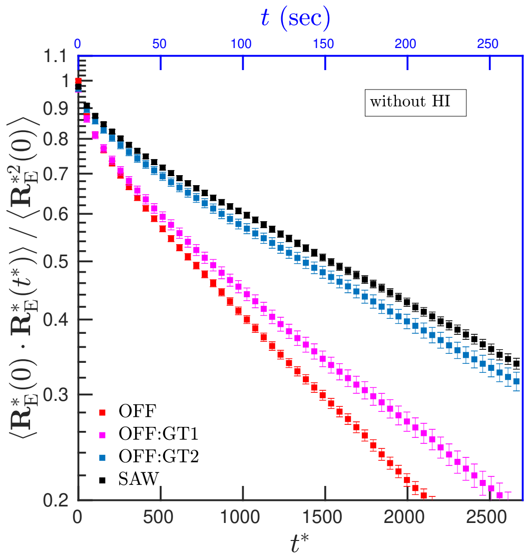


FIG. S13: Exponential decay of end-to-end auto-correlation function with time. Here the HI is not included in the simulation.

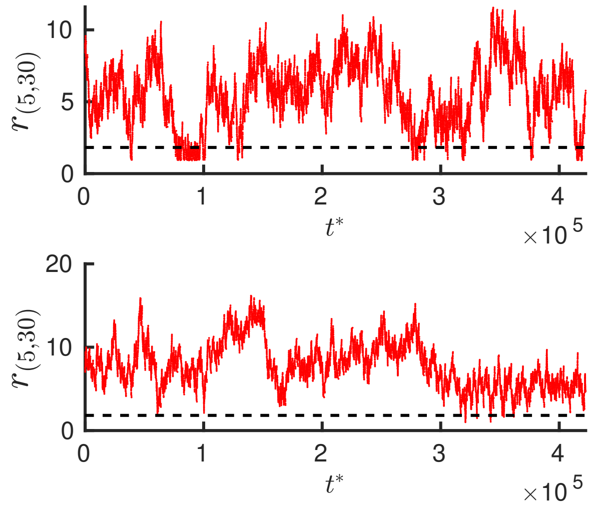


FIG. S14: Distance data from our simulation for a particular pair of beads for two randomly chosen realisation.

Matter **27**, 064115 (2015).
[20] C. M. Caragine, S. C. Haley, and A. Zidovska, Phys. Rev. Lett. **121**, 148101 (2018).
[21] T. Germier, S. Kocanova, N. Walther, A. Bancaud, H. A. Shaban, H. Sellou, A. Z. Politi, J. Ellenberg, F. Gallardo, and K. Bystricky, Biophys. J. **113**, 1383 (2017).
[22] J. Des Cloizeaux, J. Phys **41**, 223 (1980).
[23] T. Witten Jr and J. Prentis, J. Chem. Phys. **77**, 4247 (1982).
[24] B. Duplantier, J. Stat. Phys. **54**, 581 (1989).
[25] H.-P. Hsu, W. Nadler, and P. Grassberger, Macromolecules **37**, 4658 (2004).
[26] A. Santra, B. Dünweg, and J. R. Prakash, arXiv:2008.05641 (????).
[27] W. Kuhn, Kolloid Z. **68**, 2 (1934).
[28] K. Šolc, J. Chem. Phys. **55**, 335 (1971).
[29] G. Zifferer, Macromol. Theory Simul. **8**, 433 (1999).
[30] C. Haber, S. A. Ruiz, and D. Wirtz, PNAS **97**, 10792

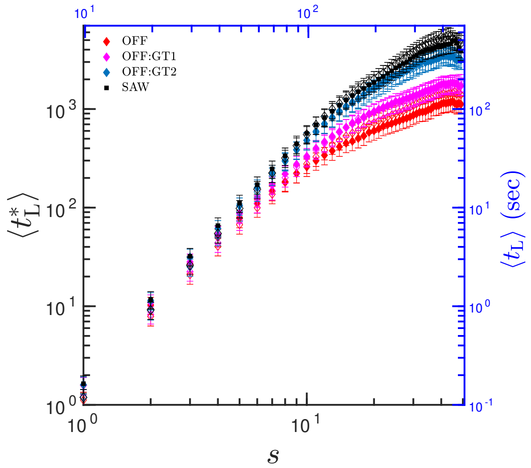


(a)

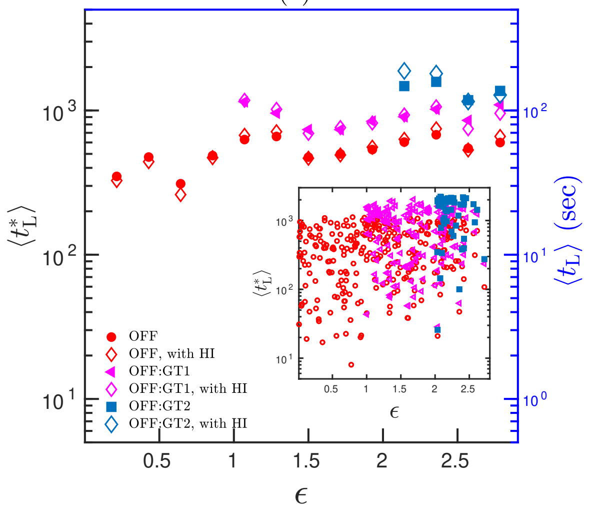


(b)

FIG. S15: (a) $\langle t^*_{\rm L} \rangle$ has a power law scaling with genomic length ($\langle t^*_{\rm L} \rangle \sim s^{\mu}$). Simulations with HI (empty symbols) and without HI (filled symbols) falls on top of each other and are indistinguishable. (b) $\langle t^*_{\rm L} \rangle$ binned and averaged over all bead pairs having same $\epsilon$ showing minimal influence of $\epsilon$. Inset: $\langle t^*_{\rm L} \rangle$ as a function of $\epsilon$ with each point representing a bead pair.

(2000).

[31] D. N. Theodorou and U. W. Suter, Macromolecules **18**, 1206 (1985).

[32] M. Bishop and J. P. J. Michels, J. Chem. Phys. **85**, 5961 (1986).

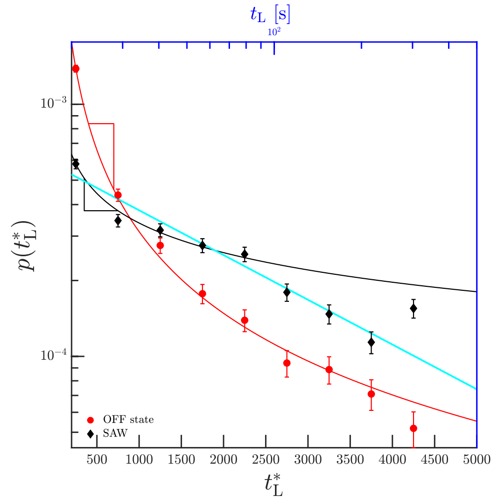


(a)

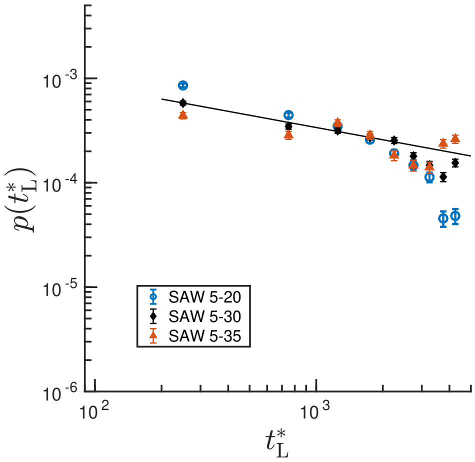


(b)

FIG. S16: (a)Fitting power-law and exponential function to examine the nature of distribution function for loop formation time ($t^*_L$) for a specific bead-pair (bead 5 and bead 30) in SAW and OFF state. A power-law function fits to the early part of SAW and the exponential function fits better to the late part. This is expected because, due to the finite size of the system, powerlaw cannot extend indefinitely; For late times, it should decay quickly as the typical time for two beads, $L$ length away, to loop cannot be too large. This is a typical finite size effect. (b)Probability distribution of loop formation time showing the effect of finite chain size in SAW. Smaller segment length such as 5-20 deviates from the power law (black solid line) much sooner compared to the larger segment lengths like 5-35.
